# Collective Excitations in α-helical Protein Structures Interacting with the Water Environment

**DOI:** 10.1101/457580

**Authors:** Vasiliy N. Kadantsev, Alexey Goltsov

**Author notes:** Corresponding authors: Alexey Goltsov.

## Abstract

Low-frequency vibrational excitations of protein macromolecules in the terahertz frequency region are suggested to contribute to many biological processes such as enzymatic catalysis, intra-protein energy/charge transport, recognition, and allostery. To explain high effectiveness of these processes, two possible mechanisms of the long-lived excitation were proposed by H. Fröhlich and A.S. Davydov which relate to either vibrational modes or solitary waves, respectively. In this paper, we developed a quantum dynamic model of vibrational excitation in α-helical proteins interacting with the aqueous environment. In the model, we distinguished three coupled subsystems, i.e. (i) a chain of hydrogen-bonded peptide groups (PGs), interacting with (ii) the subsystem of the side-chain residuals which in turn interacts with (iii) the environment, surrounding water responsible for dissipation and fluctuation in the system. It was shown that the equation of motion for phonon variables of the PG chain can be transformed to nonlinear Schrodinger equation which admits bifurcation into the solution corresponding to the weak damped vibrational modes (Fröhlich-type regime) and Davydov solitons. A bifurcation parameter is derived through the strength of phonon-phonon interaction between the side-chains and hydration-shell water molecules. As shown, the energy of these excited states is pumped through the interaction of the side-chains with fluctuating water environment of the proteins. The suggested mechanism of the collective vibrational mode excitation is discussed in connection with the recent experiments on the long-lived collective protein excitations in the terahertz frequency region and vibrational energy transport pathways in proteins.

## 1. Introduction

The concept of proteins as active dynamic systems integrates experimental and theoretical investigation of the interrelation between protein structure, dynamics, functions, and interaction with cellular environment. Currently, intense efforts are undertaken to realise the contribution of internal protein motion to protein functions such as enzymatic activity, charge/energy transport and allosteric regulation of proteins (Niessen *et al*., 2017a), (Lindorff-Larsen *et al*., 2005), (Kamerlin and Warshel, 2010). A study of the protein interaction with aqueous environment showed the link between the internal protein motion and the fluctuating of the hydration-shell water molecules and suggested an impact of this interaction on the internal protein dynamics and protein functioning (Agarwal, 2006). Protein vibration excitations spread over a wide frequency range include high-frequency localised vibrations and rotation of atom groups (e.g. C=O and N-H stretching) and the large-scale vibrations of subdomains of the proteins (Xie, Yao and Ying, 2014), (Balu *et al*., 2008).

Different types of the collective modes are reported to be excited in protein molecules, i.e. normal vibration modes (Acbas *et al*., 2014; Turton *et al*., 2014), sound waves (phonons) (Liu *et al*., 2008), and coherent vibrational states (Del Giudice *et al*., 1986; Rolczynski *et al*., 2018) which are observed on a timescale from picosecond to microsecond. Vital cellular processes are suggested to be controlled by the excitation of the long-lived collective degrees of freedom which are characterised by a weak coupling to the other ones and related to the substantially non-equilibrium processes in proteins (Mohseni *et al*., 2014). Investigation of weakly relaxing excitations discribed by the quaiparticle dynamics such as phonons, exitons, polarons and others have given an insight into a vast amount of experimental facts in solid and soft matter physics (Venema *et al*., 2016). Similarity of the non-linear mechanisms underlying formation of the collective excitations in different molecular systems suggestes that long-lived non-equilibrium states can be realised in the living systems and the concept of quasiparticles plays a crucial role in effectiveness of energy transformation, transport and storage in cells (Lambert *et al*., 2013), (Davydov, 1985). The concept of quasiparticles in biophysics was developed by Davydov and Frohlich to explain effective energy and cardge transprt in biosystems. Investigation of the physical mechanisms of the formation and physiological functions of the coherent exitations in cellular molecular structures are one of the formidable challenges in molecular biophysics and the subject of intensive experimental and theoretical works over the past decades (Engel *et al*., 2007; Acbas *et al*., 2014; Turton *et al*., 2014; Goncharuk *et al*., 2017; Niessen *et al*., 2017b), (Kuramochi *et al*., 2019).

The physical mechanism of the collective coherent excitations have been proposed and investigated early by H. Fröhlich (Fröhlich, 1968a). He has developed a phenomenological kinetic model of collective longitudinal vibrational-polar modes (phonons) excited in the 0.1 GHz - 1 THz frequency range and suggested their role in biological processes such as enzymatic catalysis, biomembrane function, protein-protein interaction, and the interaction of biosystems with microwave radiation (Fröhlich, 1970, 1980).

The general idea underlying the suggested mechanism of long-lived coherent vibrations in biosystems is in an assumption that the vibrational modes are capable of condensation into the lowest-frequency vibrational mode like the phenomenon of Bose-Einstein condensation. In contrast to Bose-Einstein condensation occurring in thermal equilibrium, Fröhlich condensation takes place in non-equilibrium condition at the energy supply and dissipation in non-linear molecular structures (Mesquita, Vasconcellos and Luzzi, 2004). Thus, excitation of the Fröhlich mode, in the form of the coherent dynamic structure, can be considered as the emergence of a space-temporal dissipative structure in accordance with Prigogine’s theory (Prigogine and Lefever, 1973) which is governed by the self-organization principals of Haken’s synergetics (Haken, 1983). At present, further investigation of the Fröhlich condensation in quantum dynamics and quasi-classical approaches were undertaken in order to determine the physical conditions of phonon condensation in proteins functioning far from thermal equilibrium (Vasconcellos *et al*., 2012), (Salari *et al*., 2011), (Preto, 2017), (Reimers *et al*., 2009), (Nardecchia *et al*., 2018).

Another type of coherent excitations in the form of solitary waves in α-helical protein structurews has been proposed theoretically by A.S. Davydov in order to describe highly effective long-distance transport of energy/charge within macromolecules (Davydov, 1985). As a result of a series of his works, the theory of soliton transport of energy/charge in the α-helical proteins has been developed (Davydov, 1977, 1985). Additionally, it was concluded that the α-helical peptide structure plays a significant role in the formation of the soliton, travelling a long distance with weak decay.

The soliton model was applied to describe the transport of energy, released in the hydrolysis of ATP and localised in the amide-I vibration (C=O bond), along the chain of peptide groups (PGs) at room temperature (Davydov, 1985). Mechanisms of energy transport of are defined in the model as the non-linear interaction of the high-frequency amide-I excitation (1667 cm^-1^) and low-frequency acoustic modes in the α-helical protein structure.

Various theoretical aspects of soliton dynamics in α-helical proteins, including soliton stability, thermalisation, solitons’ interaction and others were investigated in various approximations (Takeno, 1984), (Lawrence *et al*., 1987), (Lupichev, Savin and Kadantsev, 2015). Dadydov model was further extended to the multisoliton solutions in discrete nonlinear Schrodinger equation of amide-I excitation dynamics in the PG chain (Tchinang Tchameu, Togueu Motcheyo and Tchawoua, 2014), (Issa *et al*., 2018). An exact quantum dynamical description of intra amide group vibration transfer coupled with the low frequency longitudinal vibrations in an α-helical polypeptide was developed in the multi configuration time-dependent Hartree (MCTDH) (Tsivlin and May, 2007) and the mixed quantum–classical Liouville (MQCL) equation approximations (Freedman and Hanna, 2016). The unified theoretical approaches to the description of both Davydov soliton mode and Fröhlich condensation mode excitations at the conditions far away from thermal equilibrium in proteins were developed (Del Giudice *et al*., 1986; Bolterauer and Tuszyński, 1989; Bolterauer, Tuszyński and Satarić, 1991; Mesquita, Vasconcellos and Luzzi, 2004).

Searching for the experimental observation both of these types of coherent excitations in proteins remains an area of intense experimental research and source of lively debate (Reimers *et al*., 2009; Salari *et al*., 2011; Preto, 2017), (Nardecchia *et al*., 2018), (Weightman, 2014), (Austin *et al*., 2009). Experimental investigation of the coherent vibrational states in proteins was carried out by the IR, Raman, terahertz spectroscopy and their combinations (Nardecchia *et al*., 2018), (Lundholm *et al*., 2015) and included analysis of the possible resonance effects of microwave and terahertz irradiation on cellular functions predicted by Fröhlich (Pokorný, 1999), (Foletti *et al*., 2013), (Markov, 2015), (Fröhlich, 1980). Theoretical investigation in this field of electromagnetic biology also aims at the identification of biological structures enabling to maintain collective vibration states (α-helical protein structure, cytoskeleton microtubules, biomembranes and others) and calculation of the vibration spectrum of collective excitations containing specific frequency domains to inform an experimental search for the resonance effects of terahertz irradiation on cells (Fröhlich, 1988), (Pokorný, 2004), (Kadantsev and Savin, 1997), (Nardecchia *et al*., 2018). The resonance interaction of cells with terahertz irradiation is considered as the possible biophysical mechanism which can be explored in the development of medical applications for the non-invasive diagnostics and therapy (Foletti *et al*., 2013), (Markov, 2015), (Siegel, 2004), (Foletti *et al*., 2013).

Another type of the protein interaction with environmental factors relates to the interaction with solvent surrounding native proteins. The surface interaction of proteins with the fluctuating hydration-shell and bulk water molecules has been shown to contribute significantly to their internal vibrational dynamics and functional properties (Bellissent-Funel *et al*., 2016), (Doster, 2010).

In this work, we develop further the molecular models of the collective excitations in α-helical peptide macromolecules (Kadantsev, Lupichov and Savin, 1987; Kadantsev, Lupichcv and Savin, 1994; Lupichev, Savin and Kadantsev, 2015), (Kadantsev and Goltsov, 2018) and extend quantum dynamics approach to take into account protein interaction within the aqueous environment. In the model development, we distinguished three subsystems i.e. (i) a chain of hydrogen-bonded peptide groups (PGs) interacting with (ii) the subsystem of the side-chain residuals which in turn interact with (iii) surrounding water molecules though the phonon-phonon mechanism. We showed that consideration of vibrational coupling between macromolecules and their water environment significantly changes dynamic behaviour of the α-helical protein structures and contributes to the formation of coherent vibrational states in the PG chain. We used the quantum dynamics approach which have been developed earlier in the modeling of autolocalized states (polaron) in the intramolecular excitations of the PG chain (Kadantsev, Lupichov and Savin, 1987; Kadantsev, Lupichcv and Savin, 1994; Lupichev, Savin and Kadantsev, 2015). The developed model is used to investigate conditions when the molecular interaction between the PGs, side chains and water environment leads to the excitations of vibrational and soliton types in α-helical protein structures.

## 2. Phonons in a one-dimensional chain of hydrogen-bonded peptide groups interacting with the side-chains

The secondary protein structure, α-helix, is formed as a result of folding up of polypeptide chain in a helix due to the interaction of amino acid residues (Fig. 1a). This interaction determines space periodicity of the secondary structure in proteins, and in turn, its stability is ensured by hydrogen bonds between NH- and CO-groups of the backbone (Fig. 1b). The α-helix has a form of a coil, and its inner part consists of a tightly twisted backbone with amino acid residues directed outwards (Fig. 1c). Electrical charge distribution in the peptide group forms its electrical dipole moment equal to 3.5 D and directed along H-bond (Davydov, 1977).

**Fig. 1.**
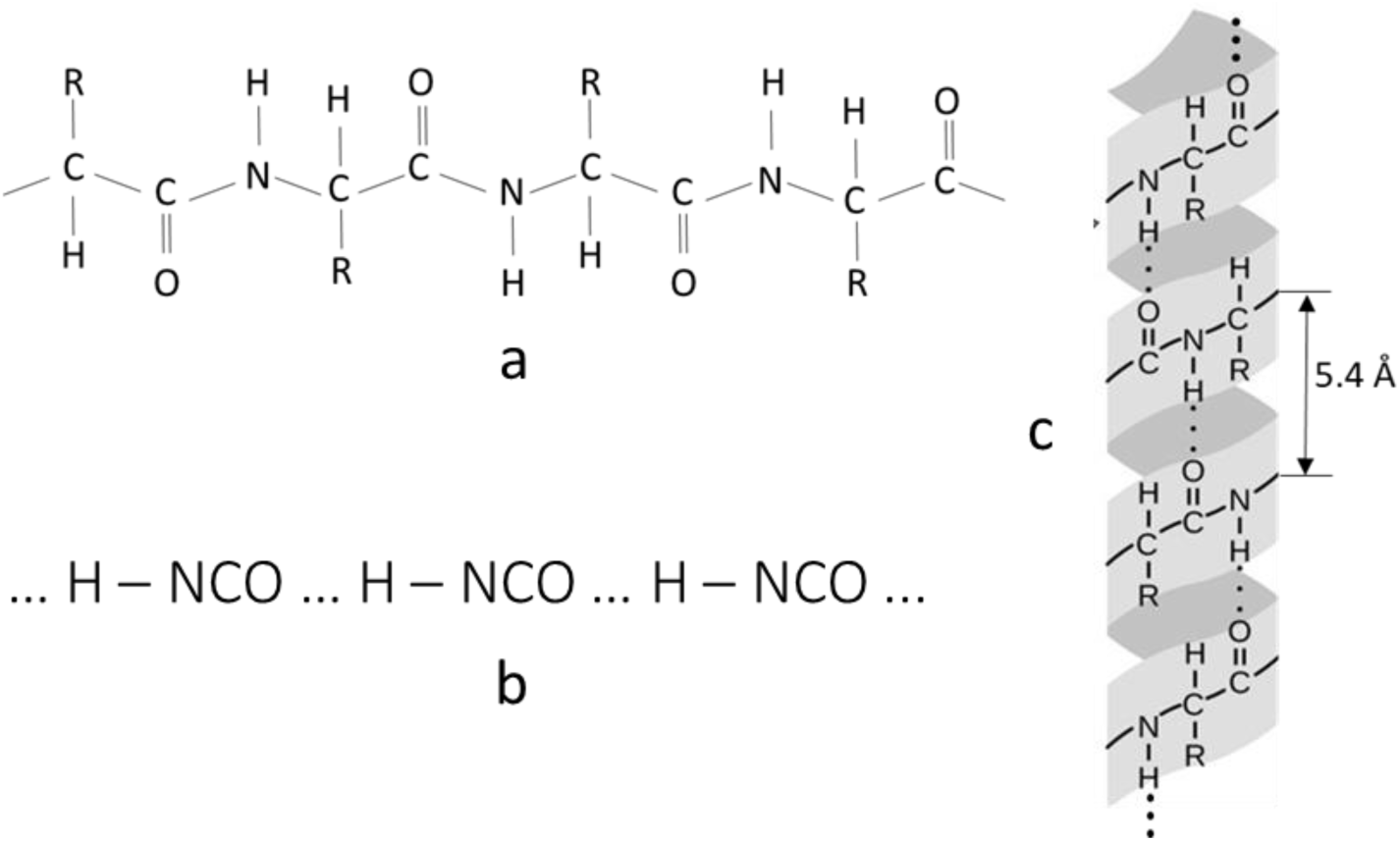
(a) A polypeptide chain with amino acid residues (R). (b) The chain of hydrogen-bonded peptide groups. (c) Structure of α-helical protein structure, where one of the three hydrogen-bonded PG chains with the side-chains of amino acid residues are shown. Hydrogen-bonds are depicted by dots.

In the model of the α-helical protein structure we considered a one-dimensional chain of hydrogen-bonded peptide groups with hydrogen bonds between (NH)- and CO-groups (Fig. 1b). We also assumed intrinsic motion of proton and the rest of the PGs (NCO) and defined equilibrium positions of the PGs in the *l*-site z=*la* (*l*=0, 1, 2, …) along the α-helix backbone (the z-axis), where *a=*5.4 Å is chain spacing. We denoted displacement of the atoms from equilibrium positions of the PG in the *l*-site by ξ_*l*,1_ for hydrogen and ξ_*l*,2_ for the rest of the PG atoms. In harmonic approximation, potential energy of the interaction between the nearest PGs and nearest chain-side of residues R is expressed by a quadratic form:

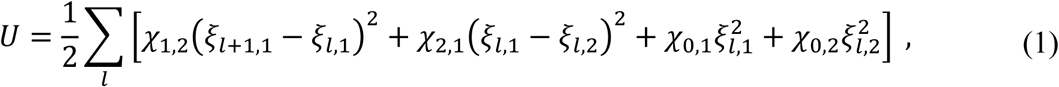

where χ_1,2_ and χ_2,1_ are elastic coefficients of hydrogen and valence bonds respectively; *χ*_0,1_ and χ_0,2_ are elastic coefficients of the interaction of protons and NCO group with the side chain of residues, respectively. In the model, we chose the following cyclic boundary condition for the PG chain

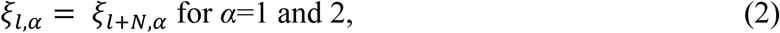

where *N* is a number of GPs in the chain.

To write the Hamiltonian of the PG chain, the operators of atom displacement from equilibrium positions were expressed through the operators of the creation 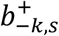 and annihilation *b*_*k,s*_ of phonons as

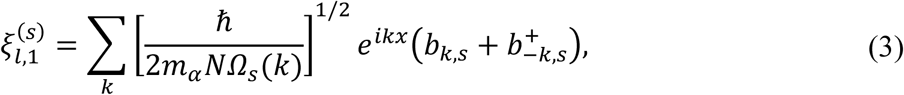

where the operators of the creation and annihilation satisfy the commutative relationships

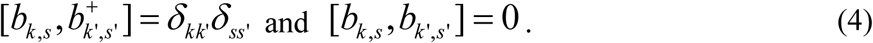

Here δ is the Kronecker symbol, *k* is the wave number taking *N* values in the first Brillouin zone

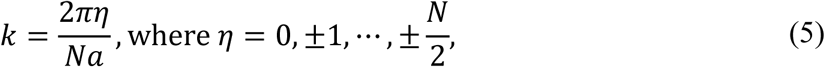

and index *s* points to either acoustic (*s*=1) or optic (*s*=2) phonons.

The Hamiltonian of phonons in the PG chain can be written in the harmonic approximation as

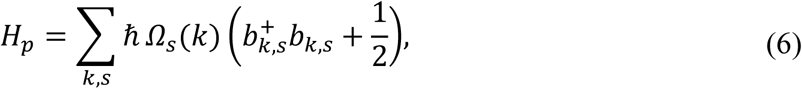

where two function *Ω*_*s*_(*k*) defines dispersion relationship for frequencies of acoustic and optic branches of the vibration in the PG chain with *s* = 1 and 2 respectively.

The dispersion relationship *Ω*_*s*_(*k*) for the PG chain with interaction defined by eq. (1) can be obtained in the form

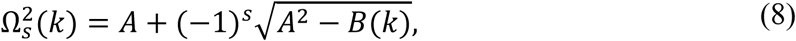

where

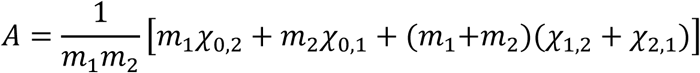

and

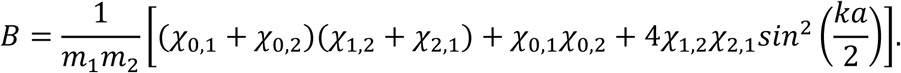

Here *m*_1_ = *m*_*p*_ and *m*_2_ = 41.7*m*_*p*_ are mass of proton and the PG group respectively. The dispersion relation (8) can be derived using the methods applied to lattice oscillation of a diatomic linear chain (Animalu, 1977). Dispersion curves calculated at the values of elastic constants of the PG chain (Lupichev, Savin and Kadantsev, 2015) are shown in Fig. 2. As seen according to dispersion equation (8), the PG chain with interaction (1) obeys a narrow band of the normal modes of vibrations *Ω*_*s*_(*k*) in the terahertz frequency range.

**Fig. 2.**
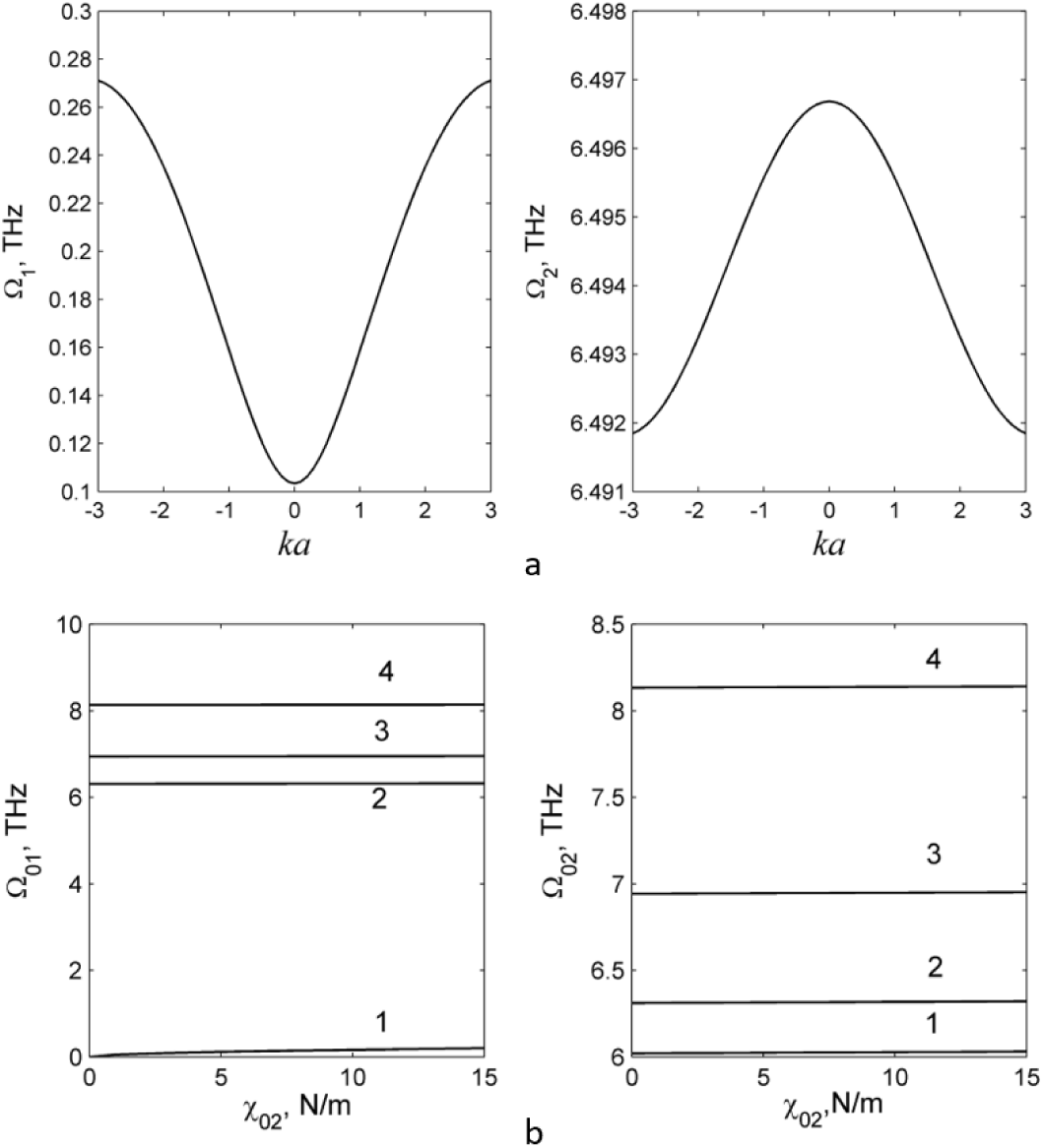
(a) Dispersion curves of acoustic (s=1, left plot) and optic (s=2, right plot) branches for the chain of hydrogen-bounded PGs calculated at the following elastic constants (Lupichev, Savin and Kadantsev, 2015): χ_1,2_ = 13.0 N*m*^−1^, χ_2,1_ = 17.0 N*m*^−1^, χ_0,1_ = 5.0 N*m*^−1^, χ_0,2_ = 0 N*m*^−1^. (b) Dependence of the frequency *Ω*_01_ (left plot) and *Ω*_02_ (right plot) (*k*=0) on constant χ_02_ at the fixed values of elastic constants χ_1,2_ = 13.0 N*m*^−1^, χ_2,1_ = 17.0 N*m*^−1^ and different constant χ_0,1_: line 1 - χ_0,1_ = 0 N*m*^−1^, line 2 - χ_0,1_ = 3.0 N*m*^−1^, and line 3 - χ_0,1_ = 25.0 N*m*^−1^.

Similarly, we introduced the phonon Hamiltonian for the side-chains of the residues which can be represented as a system of *N* oscillators of mass *M* having own frequencies of oscillation *Ω*_*t*_(*q*)

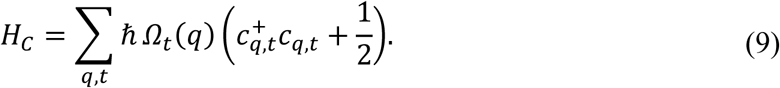

Here 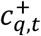 and *c*_*q,t*_ are the operators of the creation and annihilation of phonons in the side-chains which correspond to the displacement of the residues from their equilibrium positions *ζ*_*l*_(*t*). *Ω*_*t*_(*q*) are the phonon frequency of type *t* in the side-chains and *q* is the wave number. The representation of the side-chain dynamics as a set of oscillators is based on experimental data on the amino acid residue dynamics obtained by the X-ray diffraction, NMR methods and spin labelling EPR spectroscopy that provides detailed information on the side-chain order parameter and their mobility on the nanosecond timescale (Columbus and Hubbell, 2002), (Go, Noguti and Nishikawa, 1983), (Sivaramakrishnan *et al*., 2008). Moreover, molecular dynamic and normal mode calculations of protein internal dynamics showed that harmonic approximation for the modelling of the side-chain dynamics is the satisfactory approximation that gives the consistent description of the normal low-frequency modes in the range of 120-200 cm^-1^ (Go, Noguti and Nishikawa, 1983). The analysis of the complex EPR spectrum of a spin labeled side chain in α-helixes showed that the side-chain dynamics is modulated by the interaction with the backbone as well as environment (Columbus *et al*., 2001). Below we consider the interaction of the phonons in the backbone and residue side-chains in the α-helical protein, and then we include the interaction of the side-chains with aqueous environment of the proteins.

To consider interaction of the PG chain and residue side-chains in the α-helical proteins, we added the following anharmonic operator of phonon-phonon interaction to the Hamiltonian of non-interacted chains

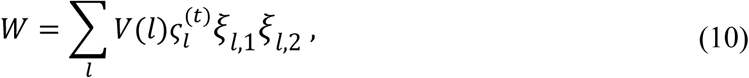

where

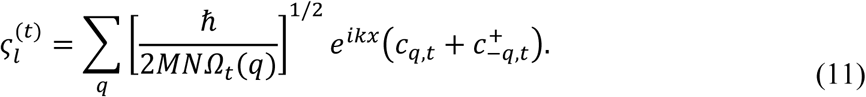

Experimental data on the α-helix backbone and side-chains interaction was derived from the analysis the EPR spectra of a spin labeled side chain that showed the tight coupling between backbone and side-chain dynamics in α-helical protein structures (Columbus *et al*., 2001)(Columbus and Hubbell, 2002).

Substitution of eqs. (3) and (11) into eq. (10) gives the operator of anharmonic perturbation in the form

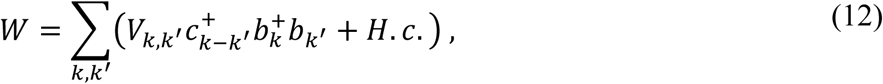

where

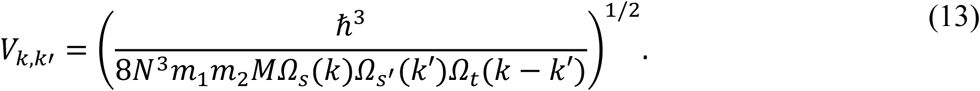

The matrix elements (13), which are different from zero on the functions of occupation numbers, corresponds to the processes occurring with energy and momentum conservation without consideration of umklapp process:

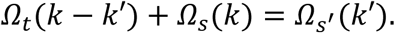

## 3. Interaction of α-helical protein structure with water environment

In the model, we considered the interaction of the α-helical protein with its environment, surrounding water molecules which form a heat reservoir with a non-zero temperature *T*. The protein-water interaction defines dissipation processes in the system and contributes to the structure and functions of proteins (Bellissent-Funel *et al*., 2016), (Kurian *et al*., 2018a), (Agarwal, 2006). In the model, we represented dynamics of the surrounding water molecules by a large set of harmonic oscillators and suggested that this system possesses collective excitation modes with dynamics like that of an oscillators’ ensemble. This approximation can be justified by experimental observation of the collective vibrational subpicosecond dynamics (phonons) of the hydrogen bond network of water molecules in hydration shells of proteins in terahertz and IR spectra (Conti Nibali and Havenith, 2014).

Note that we considered the α-helical protein interaction with water molecules only through the amino acid residues and omitted hydration of the backbone carbonyl (–C=O) and amide (–N–H) groups in the model. This approximation is reasonable in case of large charged side chains of amino acid residues which shield backbone hydrogen bonds from water molecules and stabilise α-helix protein structure in aqueous environment (Ghosh, Garde and García, 2003). Furthermore, salt-bridge formation between amino acid residues additionally stabilises α-helix and prevents it from folding. A chain of the salt-bridge can also facilitate the excitation of vibration modes along these chains as discussed in the previous section. Additionally, in the model, we neglected different dynamic properties of the hydration shells and bulk water molecules of proteins.

Then, the energy operator of the heat reservoir (surrounding water) can be written as a sum of energy operators for the independent oscillators through the operators of the creation 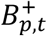 and annihilation *B*_*p,t*_ of phonons with the wave number *p* and frequencies dispersion relation *ω*_*t*_(*p*)

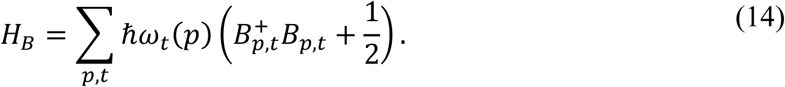

To calculate energy, *W*_*B*_, of the joint action of heat reservoir’s oscillators on the protein macromolecule we assumed that each oscillator contributes linearly to the energy *W*_*B*_ and interacts only with the residue side-chains. So, we suggested that the PG chain interacts with the environment through amino acid residues R (Fig. 1). Then, interaction operator *W*_*B*_ can be written as

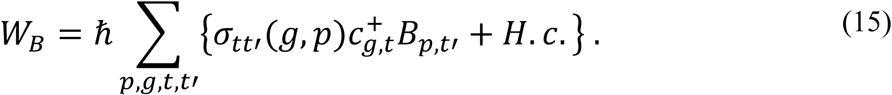

Parameter *σ*_*tt*′_(*g, p*) defines strength of the interaction between heat reservoir’s oscillators of types *t*′ and the residue side-chain oscillators of type *t*. Note that the operator (15) describes a wide class of relaxation mechanisms related to collective excitations (Lupichev, Savin and Kadantsev, 2015). The representation of the heat reservoir by a set of harmonic oscillators (Eq. 14) and protein interaction with the thermal reservoir in the form of Eq. (15) are commonly used in the modelling molecular chain dynamics particularly in the Wu-Austin Hamiltonian approximation (Columbus and Hubbell, 2002), (Vasconcellos *et al*., 2012), (Wu and Austin, 1981).

For simplicity, we dropped indexes which define phonon type, and below indexes *s, t*, and *t’* of variables mean that the specific variable belongs either to the PG chain or the side-chains or heat reservoir, respectively.

## 4. Equation of motion

One of the features of a self-organisation behaviour of complex systems is the occurrence of system instability with respect to either one or several variables (dynamic modes) when attaining a critical condition (Haken, 1983). If the rest of the modes damp the different exclusion procedures of the stable variables can be applied. As a result, the system behaviour as a whole is defined by the dynamics of a few unstable variables which governs all the damped modes. In real systems, a hierarchy of relaxation times takes place that allows applying adiabatic approximation for the exclusion of fast-relaxing variables. In the case of protein molecules, the fast-relaxing variables 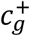 and *c*_*g*_ relate to the side-chains which directly interact with the environment (structured water). This approximation is based on the NMR experimental data on the relaxation of order parameter in backbone and side-chains dynamics (Columbus and Hubbell, 2002), (Wu and Austin, 1981). The experimental results revealed that the side-chains exhibit faster nanosecond dynamics than microsecond motion of the backbone. Moreover, solvent exposed side-chains showed more mobile and faster dynamics than those fully buried in the protein hydrophobic core.

The interaction energy of the α-helical protein with the heat reservoir *HPB* after exclusion of the variables related to the side-chain residues is defined by

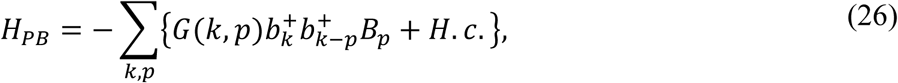

where

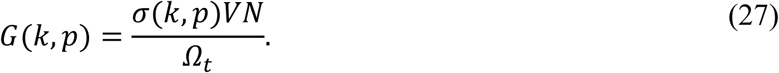

Equations of motion for dynamic variables of the PG chain and heat reservoir can be derived using Heisenberg equation (16) for the operators 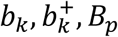, and 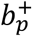, the Hamiltonian operator (21), and eqs. (22)-(27) in the forms

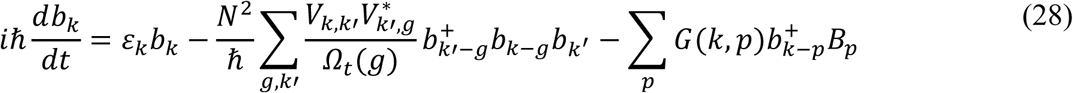

and

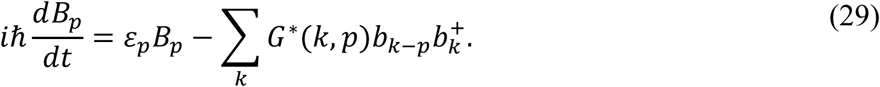

At the next step, we used the obtained eqs. (28) and (29) to derive the motion equation for the growing modes and define system dynamics near unstable stationary points. Moreover, we described dissipation in the system and express fluctuating forces which are coursed by the interaction of the protein macromolecule with the environment.

## 5. The Langevin equation for generalized coordinates of the protein macromolecule

As known, all basic (microscopic) equations of motion are invariant with respect to time reversal, that is the motion is entirely reversal, and dissipative forces, violating this invariance, cannot be expressed in the original equations. But under certain assumptions, the Langevin equations can be derived from the Heisenberg equation for a system interacting with a heat reservoir which is represented by a set of harmonic oscillators (Shibata and Hashitsume, 1978).

Heretofore, we considered the model of α-helical protein interacting with the heat reservoir and turned to classical representation of this system. Such conversion can be justified by a large number of phonons in strongly exciting modes in the protein as well as a large number of oscillators in the heat reservoir. This allowed us to represent phonon amplitudes by c-numbers and substitute operators 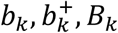, and 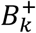 in eqs. (28) and (29) for c-numbers 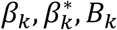, and 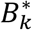 respectively. Amplitude β_*k*_(*t*) can be considered as the generalised coordinates with corresponding generalised momentum 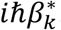. Moreover, coefficients, defining intensity of phonon interaction of the different subsystems in the model, are assumed to weakly depend on the phonon momentum. Then, eq. (29) can be integrated as the classical one that gives the solution for variable *B*_*p*_(*t*) (see details in Appendix (eqs. (A1-A7)). Substitution *B*_*p*_(*t*) (eq. A5) into eq. (28) for *b*_*k*_(*β*_*k*_) gives us the Langevin equation for phonon amplitudes *β*_*k*_(*t*) in the form

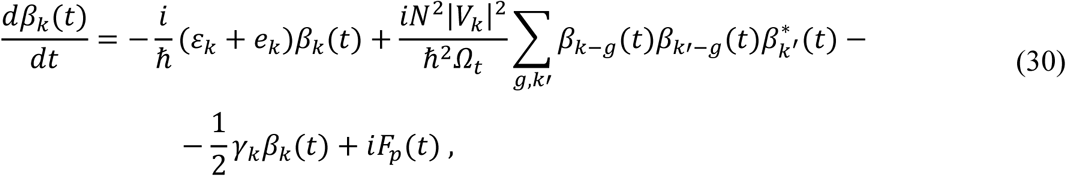

where

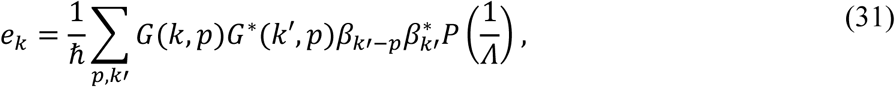

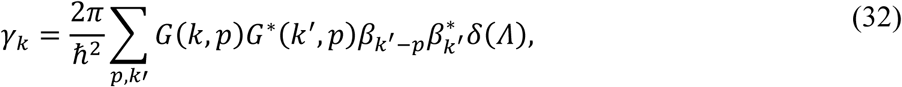

and

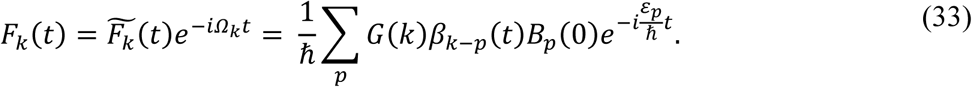

Function *F*_*k*_(*t*) (33) can be considered as a random force with the correlator (see details in Appendix, eqs. (A8-A14)):

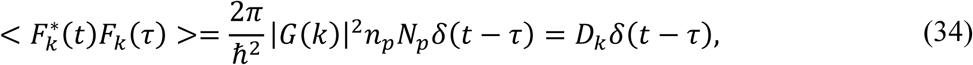

where value *D*_*k*_ can be obtained in the form (see Appendix)

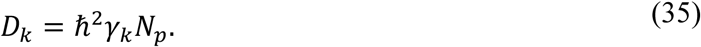

## 6. Vibrational dynamics of α-helical protein in a long wave approximation

To investigate vibrational modes in the α-helical protein at the various parameters of its interaction with the environment, we applied a long-wave approximation to derive the equation for undamped modes and investigated system dynamics in the vicinity of an unstable point. For small values *ka* ≪ 1, phonon frequencies according to eq. (8) can be written in the form

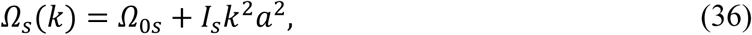

where

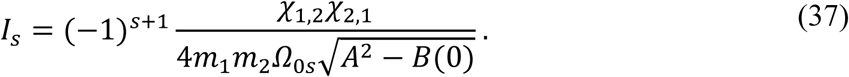

Here *Ω*_0s_ and *B* (0) are defined by eq. (8) at *k*= 0. Note that the value

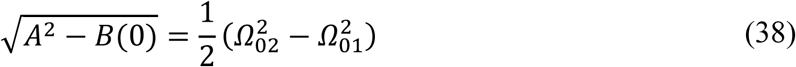

can be found from the dependence of frequencies *Ω*_0s_ on the elastic constants shown in Fig. 2. As in the long-wave approximation the value *Λ*(*k, p*) = *Λ*(*p*) does not depend on *k*, we turned to continuum limit in eq. (36) after Fourier transformation by multiplication of all terms in eq. (36) by *N*^−1/2^*e*^*ikz*^ and summing up over *k* according to eq. (36). In continuum limit, eq. (36) for photon modes with the dispersion relation (47) takes the form

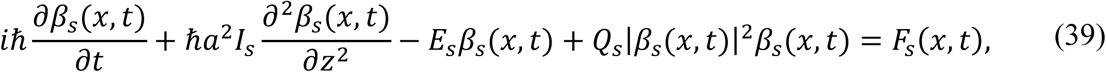

where

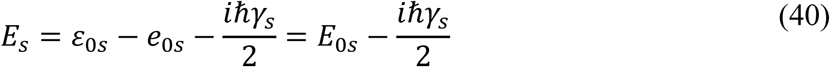

and *Q*_*s*_ is an energetic parameter of the phonon-phonon interaction between water molecules surrounding the protein and the side-chains of amino acid residues:

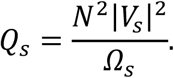

A solution of eq. (39.50) can be represented in the form

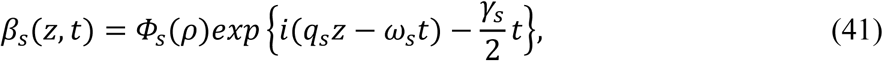

where *ρ* = *z* − *z*_0_ − *V*_*s*_, *V*_*s*_ is the velocity of excitation motion along the PG chain, and the real amplitude Φ_*s*_(ρ) satisfies the following normalization condition

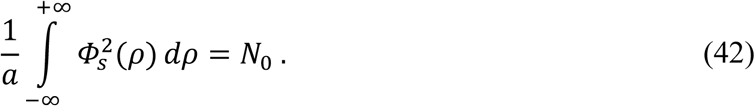

Then we considered solutions of eq. (39) at weak damping, i.e. when γ_*s*_ ≈ 0. The reasonable mechanism of the reduced relaxation of protein collective motion may be linked to the interaction of macromolecules with their environment, ordered water clusters possessing slow dynamics (Xie, van der Meer and Austin, 2002; Squire *et al*., 2013). Another mechanism of slow relaxation of the collective modes was suggested to be based on the experimental observation of vibrational wave packets of a long lifetime over 500 picoseconds in bacteriorhodopsin exposed by picosecond far IR sources (Xie, van der Meer and Austin, 2002). Authors discussed a possible mechanism of slow relaxation due to quantum effects of restricted interaction of the low-frequency collective modes with solvent and suggested a link between undamped collective vibration and the conformational transitions in proteins enriched by α-helical structures.

In this condition (γ_*s*_ ≈ 0), according to eq. (35) and fluctuation-dissipation theorem, fluctuations are small and can be neglected. Then eq. (50) takes the form:

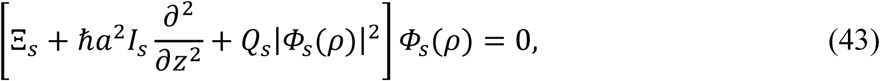

where Ξ_*s*_ is a spectral parameter connected with phonon energy by the equation:

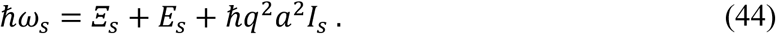

Eq. (43) has solution *Φ*_*s*_(*ρ*) = *const*:

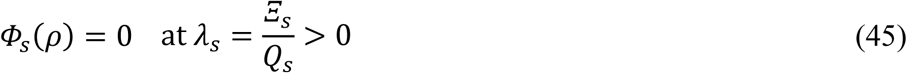

and

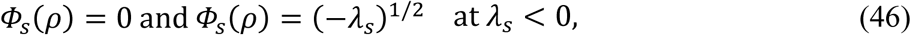

where parameter λ_*s*_ defines interaction of the PG chain with its environment. At changing λ_*s*_ the oscillation modes become unstable and their amplitudes *Φ*_*s*_(*ρ*) play a role of the order parameters of the system. Note that solutions (45) and (46) were obtained under conditions of the smallness of dissipation and fluctuations in the system. Thus, living time *τ*_*s*_ of the dynamic modes in the PG chain corresponding to nontrivial solutions (46) is less than the inverse-time of relaxation

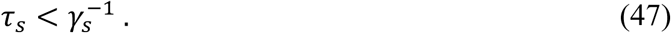

Thus, dynamics of this system is defined by weak damped (long-living) phonon modes. More detailed analysis of the dynamic equation of the type (39) is given in (Glauber, 2006), where it was shown in particular that the right-hand side of this equation can be obtained from the potential

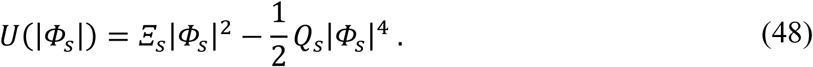

This allows writing and solving the corresponding Fokker-Planck equation and then finding a distribution function for the phonons in a coherent excitation state

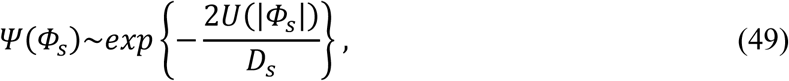

where parameter *D*_*s*_ defines intensity of fluctuating force according to eqs. (34) and (35) at *k* = 0 for both phonon branches. From eq. (49) follows that fluctuations enable the system to switch to a new state. The role of fluctuations is much significant at the transition of the system to an unstable mode at λ_*s*_ ≤ 0 when, as known, fluctuations sharply increase (Haken, 1983). At the sign change of parameter λ_*s*_, solution Φ_*s*_(*ρ*) = 0 remains one of the solutions of eq. (43). Transition of the system to the new states, corresponding to nontrivial solutions Φ_*s*_(*ρ*) ≠ 0, is possible as a result of an action of external factors including fluctuations in the environment.

Nonlinear Schrodinger equation (43) besides the solutions considered above has a solution in the form of solitary wave (soliton) travelling along the z-axis and satisfying normalisation condition (42). For any positive values of *Q*_*s*_, eq. (43) has the normalised partial solution in the form

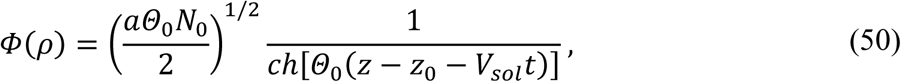

where *z*_0_ is the soliton centre, *V*_*sol*_ is soliton velocity and

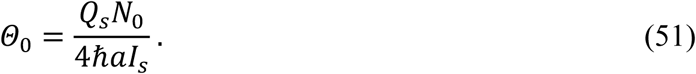

In the presence of dissipation, when γ_*s*_ ≠ 0, a solution of eq. (50) in view of eqs. (40) and (41) is written in the form:

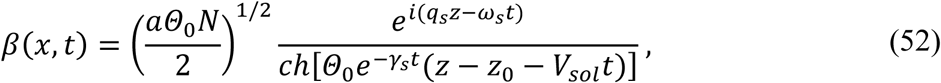

where

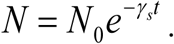

The region occupied by soliton, soliton’s width, is defined by equation

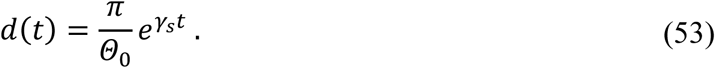

## 7. Discussion: Vibrational modes and self-organization in α-helical protein structures interacting with the environment

In the present paper, we further developed the quantum dynamic model of the collective excitations in α-helical protein structures (Davydov, 1977), (Kadantsev, Lupichov and Savin, 1987; Lupichev, Savin and Kadantsev, 2015), (Takeno, 1984). We extended this approach to taking into account the dynamics of the side-chains which is suggested to be responsible for the protein interaction within the environment and modulation of the internal dynamics of the PG chain. In the model, a side-chain-dependent coupling of the vibration excitations in the PG chain and surrounding hydrogen bond network of water molecules was considered. The model development was based on a series of approximations which are either the typical ones in the theoretical modeling of molecular chains or based on the experimental data on protein dynamics. The parameters of the model were extracted from experimental data on the structure and strength properties of peptide bonds of the α-helix.

It was showed that the equation for phonon dynamics of the PG chain interacting with the water environment can be considered as the equation for the order parameter which admits a bifurcation of its solutions. Thus, this molecular system can function in different dynamic modes, which are defined by the order parameter. Transition of a protein behavior between different states occurs at a change in the bifurcation parameter λ_*s*_ which, according to eq. (45), determines the interaction of the macromolecule with its environment. The PG chain dynamics below and above a switching threshold are significantly different. At λ_*s*_ > 0, the system is characterized by the absence of excited modes, when the PGs fluctuate with a zero-mean amplitude of the phonon modes.

At the bifurcation parameter λ_*s*_ < 0, the behaviour of the system changes so that either one or several vibrational modes become unstable and their amplitudes can grow to macroscopic values. Thus, the dynamics of the macromolecule is defined by i) the interaction of the side-chains with the fluctuation of the environment and ii) switching behaviour defined by the bifurcation parameter controlled by the protein interaction with the environment.

As shown in the model, the collective vibration modes are excited through the energy pumping to the system from fluctuating water environment (see eq. (39)). It is suggested that the vibrational energy can be channelled to the protein through the vibrational interaction of the side chains with the water molecules surrounding the protein. According to the model, the motion of the side chain residues is correlated with the motion of the hydration-shell and bulk water molecules that defined the internal dynamics of the α-helical proteins. In general, the molecular mechanisms of energy transfer from the solvent exposed side-chains on the protein surface to their internal protein motion are the subject of intense experimental and theoretical investigation (Agarwal, 2006), (Wang *et al*., 2018). As the results of this study, it has been established the vibrational energy transport pathways within enzymes which are capable of transferring required energy to the catalytic-site from dynamical fluctuations of the solvent. The mechanisms of energy pumping and transferring can be determined by the interaction of ordered water clusters bound to the hydration shell of proteins and polar amino acid residues (Goncharuk *et al*., 2017). Additionally, the hydrogen bond networks of water molecules which possess the collective vibrational sub-picosecons dynamics and propagating phonon-like modes in terahertz and IR spectra can also contribute to this mechanism (Conti Nibali and Havenith, 2014; Niessen *et al*., 2017b).

According to our model, the emergence of space-temporal structures in the form of solitary waves (Davydov regime) are possible in the PG chain. Analysis of the soliton mode showed that (i) soliton formation is governed by phonon-phonon interaction of the PGs with radical chains in α-helical proteins and (ii) the Davydov regime realises at the bifurcation parameter value corresponding to the excitation of collective vibrational modes (Fröhlich-like regime). This result showing the coexistence of the Frohlich-like vibration and Davydov soliton excitation in the same region of the bifurcation parameters agrees with similar results obtained in different approaches such as Fröhlich kinetic rate equations, Wu-Austin Hamiltonian and other methods (Del Giudice *et al*., 1986; Bolterauer and Tuszyński, 1989; Bolterauer, Tuszyński and Satarić, 1991; Mesquita, Vasconcellos and Luzzi, 2004).

The obtained results showed that instability of vibrational modes, which are induced by a change in the parameter of protein-environment interaction, can cause a formation of the new macroscopic space-temporal structures in the protein system. The joint action of random and deterministic forces leads to the switching of the system to a dynamic state, characterised by cooperative behaviour of its subsystems. i.e. the backbone, side-chains of amino acid residues, and hydrogen bond networks of surrounding water molecules.

Note, that the developed model of the α-helix interacting with its environment needs further development to build a more realistic model by considering tertiary interactions of α-helical structures in native proteins. As the α-helical structure is not stable in aqueous solution in the absence of tertiary interactions, our model in the current version can be directly applied to unfolded α-helical structures which are stable in solution. Such stable structures are the α-helical proteins and the α-helical Fs-peptide enriched with polar amino acid residues (e.g. an alanine-rich peptides) which are stable in water environment due to shielding of backbone hydrogen bonds from water molecules (Ghosh, Garde and García, 2003). These α-helical structures interact with water molecules through only amino acid residues that was considered in our model (section 3). Moreover, α-helical structures form more complex ones stable in the solution such as the coiled-coil ones, supercoils (a superhelix) as well as α-helix barrel structures due to hydrophobic interaction of the non-polar side-chains (Lupas and Bassler, 2017). The amphipathic α-helical coiled-coil structures play a significant role in molecular recognition and protein-protein interaction. For example, the leucine zipper (coiled-coil) structures are responsible for recognition and binding of the transcription factors with the DNA promoter regions of about short (∼20) nucleotide sequence (Lupas and Bassler, 2017). Excitation of the vibrational modes in the superhelices can be applied to explain the molecular recognition mechanism, long-range protein-protein (Fröhlich, 1968a) and protein-DNA interaction (Kurian *et al*., 2018b), (Christopher J. Oldfield *et al*., 2005).

Progress in terahertz spectroscopic techniques and their combination with other spectroscopic methods led to a revival of interest in the experimental observation of Fröhlich coherent excitation in biological structures (Pokorný, 2004; Weightman, 2014). Recently, a search for experimental evidence of long-range quantum coherent states in the proteins was undertaken in experimental investigation of the lysozyme protein interaction with terahertz irradiation of 0.4 THz (Lundholm *et al*., 2015). Authors observed the excitation of longitudinal compression modes of microsecond lifetime in the α-helix and attributed this underdamped low-frequency collective vibration to the Fröhlich condensation mode, excited by terahertz radiation.

Experimental investigation of the coherent vibrational dynamics in proteins was intensified by the observation of long-lived coherent excitonic states in light-harvesting proteins in photosynthetic bacteria (Engel *et al*., 2007). The results of 2D IR coherent spectroscopy suggests that the coherent vibrations in photosynthetic pigment–protein complexes contribute to the effective electron and energy transport due to the electron-vibrational couplings (Kolli *et al*., 2012; Chenu *et al*., 2013; Duan *et al*., 2017; Rolczynski *et al*., 2018). It is notable that Fröhlich in 1968 proposed the role of coherent longitudinal electric modes (polarization waves) of low frequency (0.01 THz – 1 THz) in the storage of light energy in photosynthesis (Fröhlich, 1968b).

In the recent intriguing work, the joint theoretical and experimental investigation of out-of-equilibrium collective oscillation of the Fröhlich type was carried out in the bovine serum albumin (BSA), the protein mainly composed of α-helical structures (Nardecchia *et al*., 2018). Using complementary THz spectroscopy, the strong resonance around 0.3 THz in absorption spectrum was observed, when optical pumping to the vibrational modes of the BSA took place through excitation of fluorochromes. Authors also developed a classical version of the Fröhlich model and showed that phonon condensation is possible in nonequilibrium state of the protein. Note, that the resonance absorption spectrum in the terahertz frequency region for α-helical protein structures was defined by one of the authors (VNK) in the framework of the quantum dynamics model which described the transport of electron captured by the moving acoustic soliton (electrsoliton) along the PG chain (Kadantsev and Savin, 1997).

In this work, we further developed the quantum dynamics model of the α-helical protein structure and extended it to taking into account protein interaction with the water environment. This model can be applied to the analysis of the current experimental data on the collective vibration excitations in α-helical protein structures under the different mechanisms of energy supplying and the vibrational energy transport along the quasi-linear dynamical pathways in proteins.

## Conflict of interest

The authors report no conflict of interest.

## Appendix: Exclusion of the variables related to the side-chain dynamics

Eq. (29) can be integrated as the classical one that gives the solution in the form:

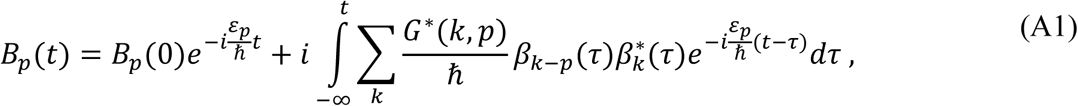

where *B*_*p*_(0) is the initial value of amplitude *B*_*p*_(*t*) at *t*=0.

Then we introduced new variable 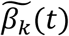

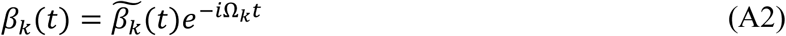

and used below the previous notation *β*_*k*_(*t*) for 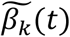. Then we applied adiabatic approximation commonly used in the modelling of cooperative systems, i.e. the relaxation times of the strong exciting phonon modes become longer in comparison with the typical relaxation times for the heat reservoir variables. This allows factoring out the preexponential term in eq. (A1) and obtain it in the form

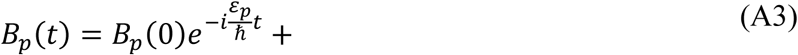

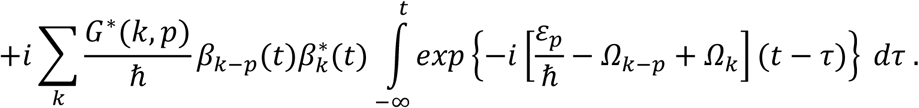

Integral in eq. (A3) gives

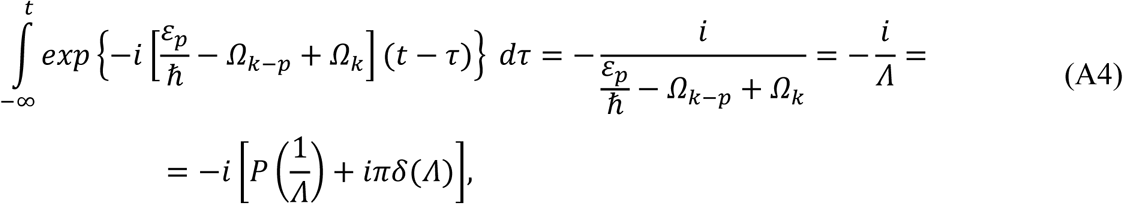

where *P* is a symbol of principal value. Finally, *B*_*p*_(*t*) is obtained in the form

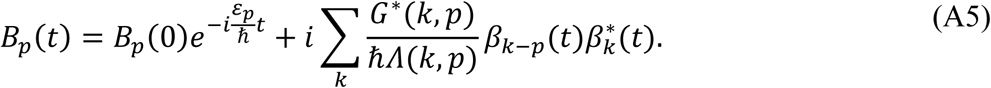

Substitution of eq. (A5) into eq. (28) gives us equation for *β*_*k*_(*t*)

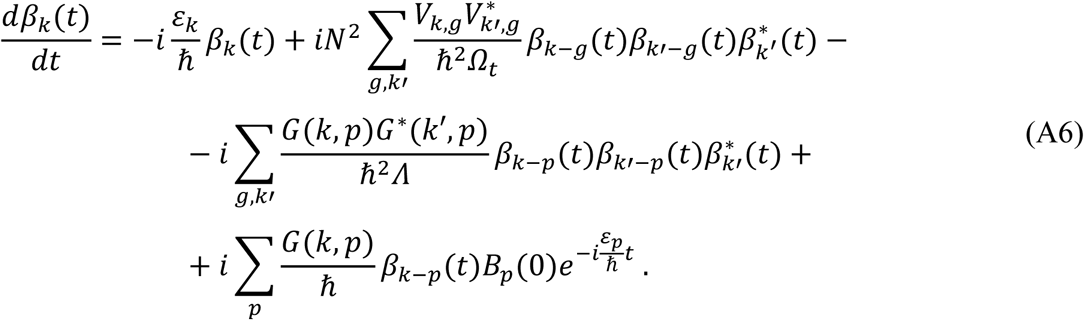

Eq. (A6) is the Langevin equation for phonon amplitudes *β*_*k*_(*t*) written in the form of eq. (30)

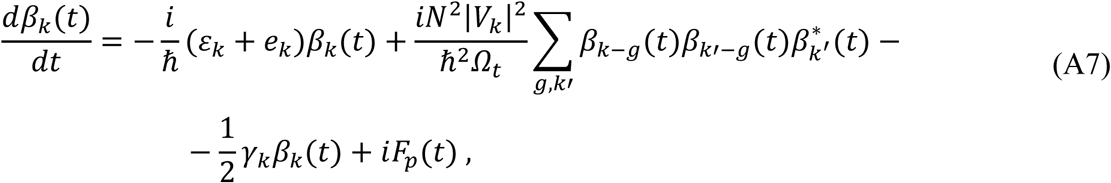

where *e*_*k*_, *γ*_*k*_ and *F*_*k*_(*t*) are defined by eqs. (31-33).

The correlator of random force *F*_*k*_(*t*) (eq. 33)

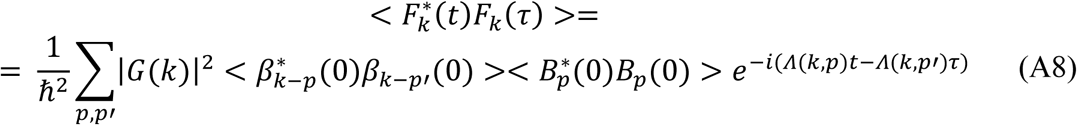

can be calculated if we assume that amplitudes *β*(0) and *B*(0) are not correlated at the initial time, i.e.

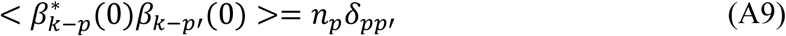

and

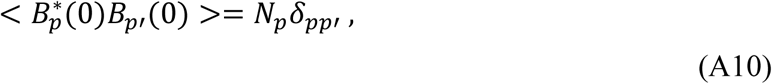

where

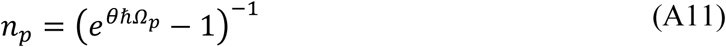

and

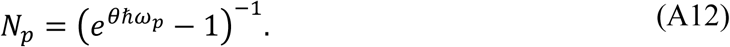

Here θ = 1/*k*_*B*_*T*.

Substitute eqs. (A9)-(A12) into eq. (A8) and take into consideration that the main contribution in a sum in eq. (A8) is given by terms *Λ*(*k, p*) = 0 at not too small values of a difference (*t* − τ). Then we finally obtained the correlator in the form of eq. (34)

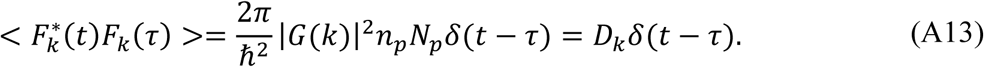

It can be shown that after averaging in eq. (32) for parameter *γ*_*k*_ with consideration of eq. (A12) and substitution of this result into eq. (A13), the value *D*_*k*_ can be written in the form of eq. (35)

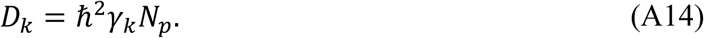

## Notes

### Competing Interest Statement

The authors have declared no competing interest.

### Summary of Updates

typos correction; reduction; Appendix added; additional text in Discussion

